# SARS-CoV-2 infection induces soluble platelet activation markers and PAI-1 in the early moderate stage of COVID-19

**DOI:** 10.1101/2021.08.23.457378

**Authors:** Abaher O. Al-Tamimi, Ayesha M. Yusuf, Manju N. Jayakumar, Abdul W. Ansari, Mona Elhassan, Fatema AbdulKarim, Meganathan Kannan, Rabih Halwani, Firdos Ahmad

## Abstract

**Introduction:** Coagulation dysfunction and thromboembolism emerge as strong comorbidity factors in severe COVID-19. However, it is unclear when particularly platelet activation markers and coagulation factors dysregulated during the pathogenesis of COVID-19. Here, we sought to assess the levels of coagulation and platelet activation markers at moderate and severe stages of COVID-19 to understand the pathogenesis.

**Methods:** To understand this, hospitalized COVID-19 patients with (severe cases that required intensive care) or without pneumonia (moderate cases) were recruited. Phenotypic and molecular characterizations were performed employing basic coagulation tests including PT, APTT, D-Dimer and TFPI. The flow cytometry-based multiplex assays were performed to assess FXI, anti-thrombin, prothrombin, fibrinogen, FXIII, P-selectin, sCD40L, plasminogen, tissue-plasminogen activator (tPA), plasminogen activator inhibitor-1 (PAI-1) and D-Dimer.

**Results:** The investigations revealed induction of plasma P-selectin and CD40 ligand (sCD40L) in moderate COVID-19 cases which were significantly abolished with the progression of COVID-19 severity. Moreover, a profound reduction in plasma tissue factor pathway inhibitor (TFPI) and FXIII were identified particularly in the severe COVID-19. Further analysis revealed fibrinogen induction in both moderate and severe patients. Interestingly, an elevated PAI-1 more prominently in moderate, and tPA particularly in severe COVID-19 cases were observed. Particularly, the levels of fibrinogen and tPA directly correlated with the severity of the disease.

**Conclusions:** In summary, induction of soluble P-selectin, sCD40L, fibrinogen and PAI-1 in moderate COVID-19 cases suggests the activation of platelets and coagulation system before patients require intensive care. These findings would help in designing better thromboprophylaxis to limit the COVID-19 severity.

## Introduction

Severe acute respiratory syndrome coronavirus-2 (SARS-CoV-2) infection often leads to respiratory conditions like acute respiratory distress syndrome (ARDS) and pneumonia in severe cases.^1-2^ Preliminary cohort studies show a higher incidence of venous thromboembolic events in COVID-19 patients who particularly required intensive care.^3-4^ Acute respiratory disease progression includes an early infection phase in which the virus attacks epithelial cells using angiotensin-converting enzyme 2 (ACE2) receptors, which leads to pneumonia and induction of severe systemic inflammation.^5^ Coagulopathy is one of the most concerning features in severe COVID-19 patients which is characterized by prolonged prothrombin and activated partial thromboplastin times, elevated D-dimer, and fibrinogen levels.^6-7^ The elevated coagulopathy markers are found to be associated with a higher mortality rate.^8-9^

SARS-CoV-2 –induced endothelial damage releases a variety of soluble markers of endothelial dysfunction including von Willebrand factor (VWF), soluble E-selectin, P-selectin, thrombomodulin.^10-11^ Such biologically active molecules eventually promote platelet activation. Platelet receptors regulate not only thrombosis but also initiate delayed cellular activities underlying inflammatory response against diverse pathogens, including viruses.^12^ Tissue factor pathway inhibitor (TFPI) is a physiological inhibitor of TF and plays a critical role in balancing the initiation phase of coagulation. TFPI is a serine protease inhibitor that inhibits TF-FVIIa and prothrombinase complex (FXa/FVa) and regulates thrombin generation.^13-14^

In a variety of infections, endothelial cells release proinflammatory cytokines such as TNF-α, IL-1, and IL-6 which in turn not only increases the expression of TF on the endothelial lining but also increases the plasminogen activator inhibitor-1 (PAI-1). Endotoxin-induced expression of TF and PAI-1 by endothelium thus may provide a stimulus that promotes thrombotic complication. ^15-16^ Several viral infections including dengue and influenza were found to be associated with platelet activation and inflammation through different molecular pathways.^17^ The inflammatory reaction can also lead to a release of reactive oxygen species (ROS) and proteases which contribute to endothelial damage and further pathological remodeling of the venous wall.^18^

Studies have revealed that patients infected with SARS-CoV-1 or MERS virus develop fibrin/thrombi within the pulmonary vasculature.^19^ Studies with SARS-CoV-1 infected mice further suggest the abnormal expression of procoagulant genes such as thrombin, VII, XI, XII, and plasminogen activators especially in those who had fatal consequences.^20-21^ Sepsis-induced coagulopathy (SIC) and disseminated intravascular coagulopathy (DIC) have been documented with severe disease, especially in non-survivors COVID-19 cases.^6^ Though the dysregulation of plasma CD40 ligand (sCD40L)^22^, TFPI^23^, tPA, PAI-1^24^ and FXIII^25^ have already been reported, most of the studies were focused particularly on severe COVID-19. However, it is still unclear particularly when these key platelet activation markers and components of the coagulation system dysregulated during the COVID-19 pathogenesis and how they correlate with the progression of disease severity.

Here we report increase levels of plasma P-selectin and CD40 ligand (sCD40L) in the early moderate stage and the levels decline with the progression of the COVID-19 severity. Fibrinolysis pathway analysis revealed a similar pattern of PAI-1, a higher level in moderate vs. severe patients, and an elevated level of tPA only in severe COVID-19 cases. These findings provide strong evidence that platelet activation induces in the early stage of SARS-CoV-2 infection when COVID-19 patients present only moderate symptoms. COVID-19 severity advances with the activation of the TF pathway, the elevation of fibrinogen, and FXIII deficiency.

## Patients and Methods

The study was conducted on COVID-19 patients, hospitalized in Rashid Hospital, Dubai between May-June 2020. SARS-CoV-2 infection was confirmed through the reverse transcription-polymerase chain reaction (RT-PCR). A total of 30 (15 moderate + 15 severe) hospitalized COVID-19 cases and 10 healthy controls (8 Males: 2 Females), age ranging from 25-43 years, were included in the study. The family history and blood samples were collected from patients and healthy controls after getting written informed consent. The ethical approval to conduct the proposed studies was taken from the institutional ethical review boards of the University of Sharjah and Dubai Health Authority.

### Inclusion and exclusion criteria

The hospitalized moderate (without pneumonia or the requirement of intensive care) and severe COVID-19 patients (presented pneumonia and/or acute respiratory distress syndrome confirmed by Chest X-ray or CT scan and, required intensive care) were included in the study. Patients with a preexisting history of cardiovascular diseases including those with myocardial infarction, stroke, or deep vein thrombosis (DVT) were excluded from the study. Patients were also excluded if received thromboprophylaxis like low molecular weight heparin or other anti-coagulant. The healthy controls were recruited after taking the medical history and informed consent. Individuals on any type of medication or with a recent history of any medical conditions were excluded. Individuals without any current medications and/or a recent history of any disease were only included as healthy control.

### Blood sample collection and plasma preparation

Blood samples from COVID-19 patients and healthy controls were collected in sodium citrate or EDTA-coated vials and processed quickly. The whole blood sample was centrifuged at 700*g* for 10 minutes and platelet-poor plasma was separated quickly and transferred to a fresh tube and stored at -80°C until further use.

### Clinical laboratory investigation

Upon hospitalization, basic laboratory investigations including complete blood count (CBC), prothrombin time (PT), activated partial thromboplastin time (APTT), D-dimer, C reactive protein (CRP) were performed. PT, APTT, and D-dimer assays were performed using semi-automated coagulometer STAGO-STAR, and complete blood count (CBC) was measured under Beckman Coulter. The level of CRP was measured using Cobas b101 system and plasma ferritin level was measured using an immunoturbidimetric assay (Cobas 6000, Roche Diagnostics).

### Flow cytometric detection of coagulation, thrombosis, and fibrinolysis markers

To detect the plasma levels of coagulation, thrombosis and fibrinolysis markers including FXI, anti-thrombin, prothrombin, fibrinogen, FXIII, P-selectin, sCD40L, plasminogen, tissue-plasminogen activator (tPA), plasminogen activator inhibitor-I (PAI-I) and D-Dimer, we performed multiplex bead-based assays using LegendPlex Human thrombosis (BioLegend #740906) and LegendPlex Human Fibrinolysis Panels (BioLegend #740761) following the manufacturer’s instruction. Briefly, diluted plasma samples from COVID-19 patients and healthy controls were used to measure the levels of soluble markers. Data were acquired under BD FACS Aria III using FACS Diva software and data analysis was done using Legendplex software (Biolegend, USA). All the samples were assessed in duplicate and the average was taken as a final reading.

### Quantification of human tissue factor pathway inhibitor

To quantify the tissue factor pathway inhibitor (TFPI) levels in the plasma samples of the COVID-19 patients and healthy controls, we performed a sandwich enzyme-linked immunosorbent assay (ELISA) using Human TFPI ELISA kit (Abcam# ab274392) according to the manufacturer’s instruction. All the samples were assessed in duplicate and the average was taken as a final reading.

### Statistics

Data group differences were evaluated for significance using unpaired *t*-test or one-way ANOVA followed by Tukey’s post-hoc test for multiple comparisons (Graph Pad Prism Software Inc., San Diego, CA). Data are expressed as mean ± SEM. For all tests, a *P*-value <0.05 was considered for statistical significance.

## Results

### Phenotypic and clinical presentation

The age range of recruited patients was between 32-69 years (28 Males: 2 Females) and the majority of the cases were of Asian background. Laboratory investigations revealed a trend of prolonged PT irrespective of the COVID-19 severity. APTT was within the normal range in the moderate and a trend of prolongation was seen in the severe group. The platelet counts were in the normal range in both patient groups. As expected, the D-dimer level both in moderate and severe groups was higher than the normal range and the level was significantly higher in the severe vs. moderate COVID-19 patients. The laboratory investigations further suggested the induction of inflammatory reaction in COVID-19 cases in which an elevated level of plasma c-reactive protein (CRP) was identified in both moderate and severe cases. Moreover, increased inflammation was further indicated by a higher number of white blood cells (WBCs) including absolute lymphocyte counts (ALC), particularly in severe COVID-19 cases. The level of plasma creatinine was unchanged and profound induction of ferritin level was seen in both patient groups (**Table**).

**Table:**
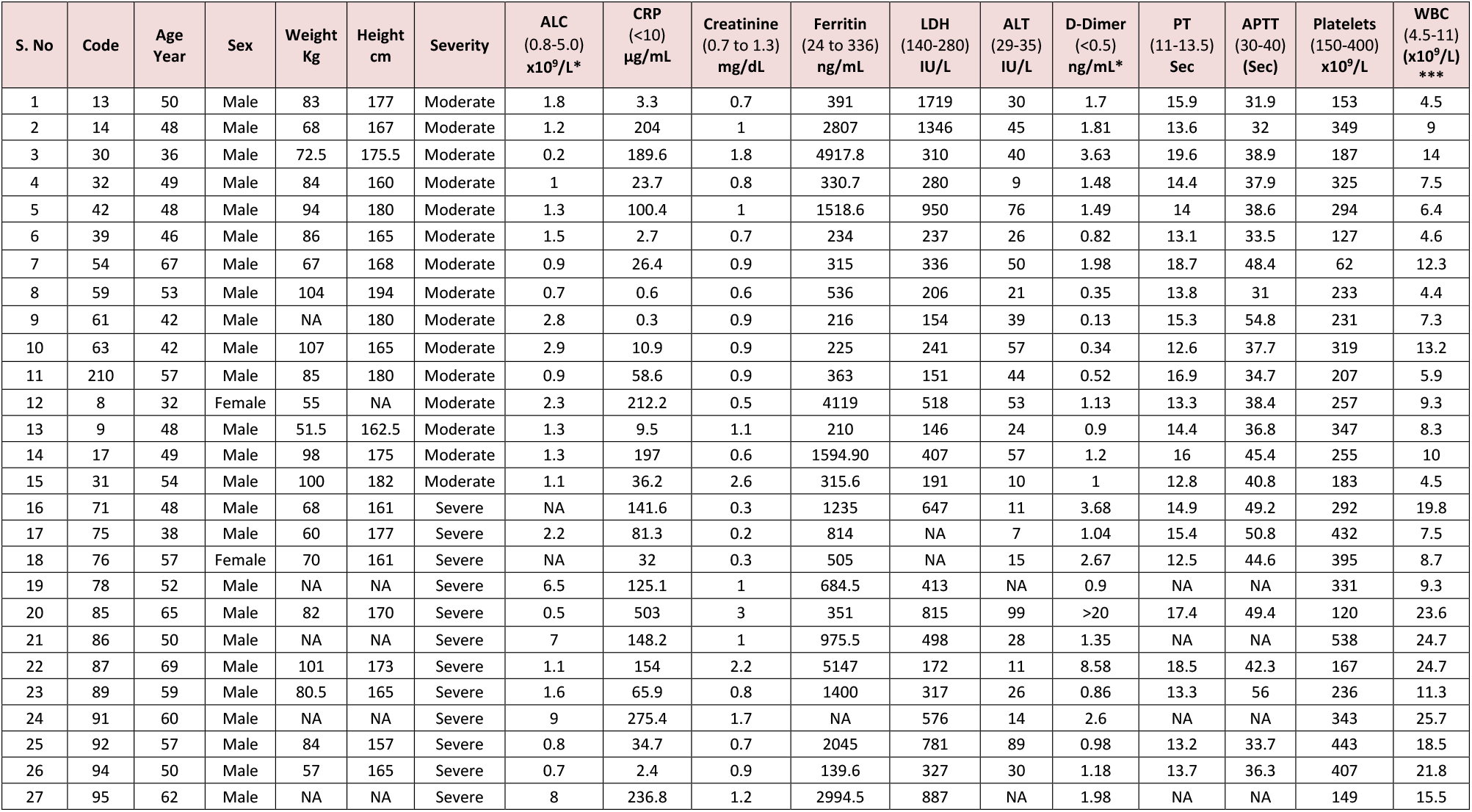
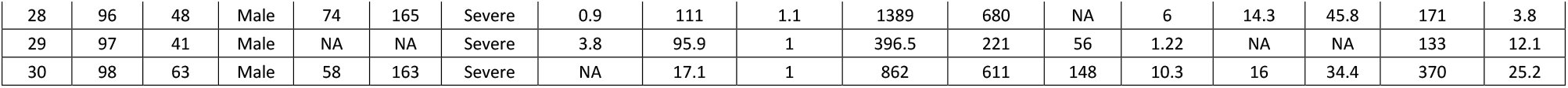
Table shows different clinical parameters of included moderate and severe COVID-19 patients. **ALC**; absolute lymphocyte count, **CRP**; C-reactive protein, **LDH**; Lactate dehydrogenase, **ALT**; Alanine transaminase, **PT**; Prothrombin time, **APTT**; Activated partial thromboplastin time; **NA**, not available. The normal ranges are provided in the bracket. *, p<0.05; ***, p<0.0005 for comparison between the moderate and severe groups.

### SARS-CoV-2 induces the release of soluble platelet activation marker

Vascular inflammation often leads to endothelium damage and platelet activation through releasing numerous intermediate biologically active molecules.^26-28^ Therefore, we sought to identify the stage of COVID-19 where the dysregulation of platelet activation markers occurs. In this series, the level of different soluble plasma markers including P-selectin and sCD40L were assessed in moderate and severe COVID-19 patients. Indeed, P-selectin was found to be markedly elevated in the moderate group which was significantly abolished in the severe COVID-19 cases (**Fig. 1A-B**). Similarly, the level of sCD40L was found significantly elevated in the moderate COVID-19 cases and the level of sCD40L profoundly diminished with the advancement of severity (**Fig. 1C**). These findings strongly suggest that SARS-CoV-2 triggers the release of P-selectin and sCD40L potentially from activated endothelial cells and induces platelet activation. The activated platelet further contributes to the elevated levels of P-selectin and sCD40L and thrombosis.

**Figure 1:**
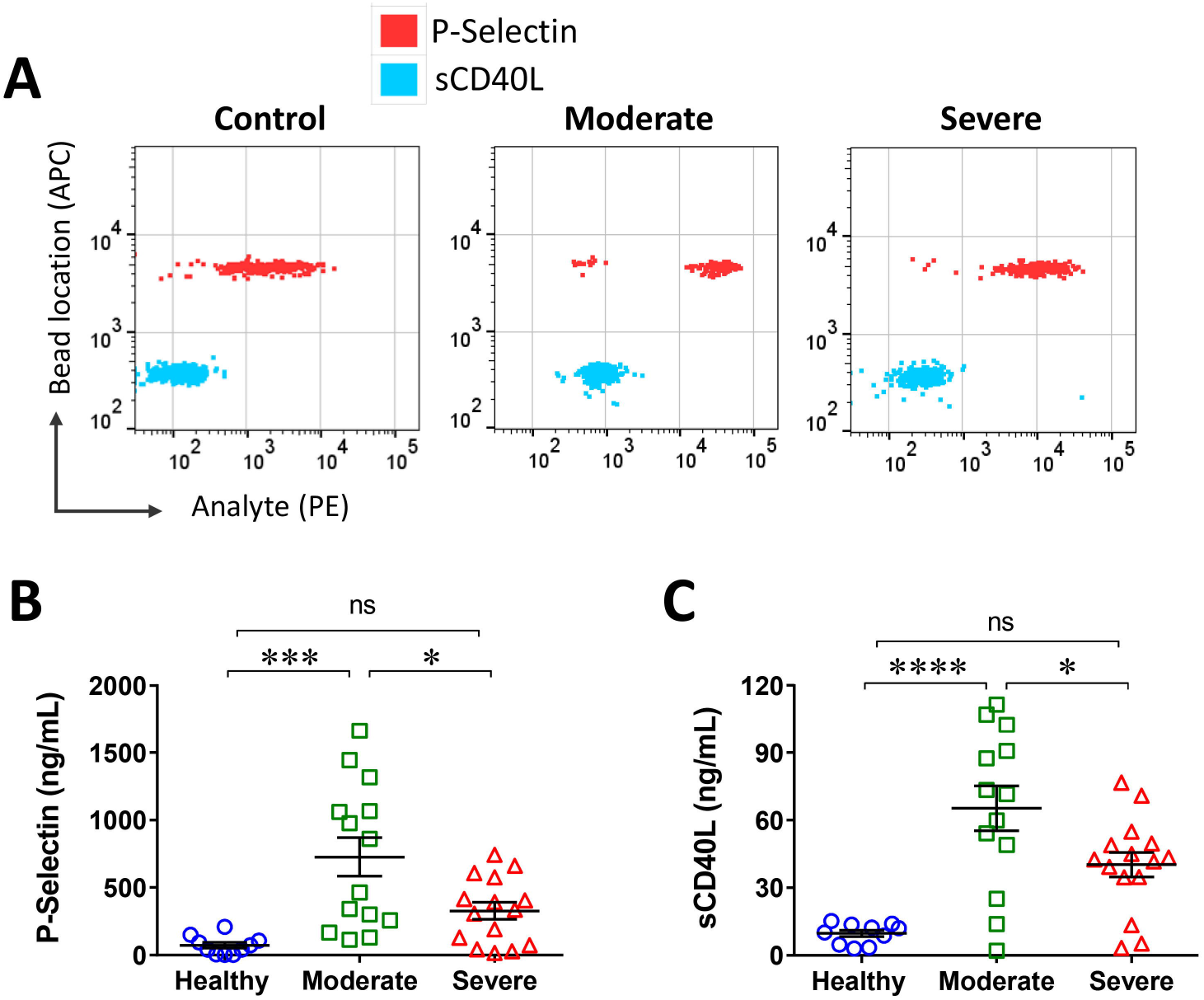
Plasma prothrombotic markers level increase at the moderate stage of COVID-19. (**A**) Representative flow cytometry scatter dot plots show the levels of P-selectin and soluble CD40 ligand (sCD40L) in moderate and severe COVID-19 patients vs. healthy controls. Scatter dot plots show significantly elevated levels of (**B**) P-selectin and (**C**) sCD40L in moderate patients and significantly lower levels in the severe vs. moderate COVID-19 patients. *, p<0.05; ***, p<0.0005; ****, p<0.0001.

### SARS-CoV-2 infection suppresses tissue factor pathway inhibitor, FXIII and induces fibrinogen

Accumulating data sets suggest that severe SARS-CoV-2 infection leads to coagulation dysfunction characterized by increased thrombin generation and elevated levels of D-Dimer ^2,6,29^. Therefore, next, we assessed the potential dysregulation of intrinsic, extrinsic, and/or common coagulation pathways that might contribute to thrombin generation and ultimately platelet activation. We observed a comparable FIX level in moderate and severe cases vs. healthy controls (**Fig. 2B**). However, extrinsic or TF pathway analysis revealed a trend of decline in plasma TF pathway inhibitor (TFPI) level in moderate patients which was significantly lower levels in the severe COVID-19 cases compared to healthy controls (**Fig. 2C**). The assessment of the common coagulation pathway suggests unchanged plasma prothrombin and antithrombin levels in both moderate and severe cases (**Fig. 2A, D&E**). Interestingly, fibrinogen level was found markedly upregulated in moderate cases which further elevated with the progression of COVID-19 severity (**Fig. 2F**). In stark contrast to fibrinogen level, a trend of lower FXIII level was observed in moderate cases, and the level further declined in severe COVID-19 (**Fig. 2G**). These findings indicate that SARS-CoV-2 -infection dysregulates the extrinsic as well as common pathways by attenuating the plasma TFPI and FXIII levels, and by upregulating the fibrinogen level.

**Figure 2:**
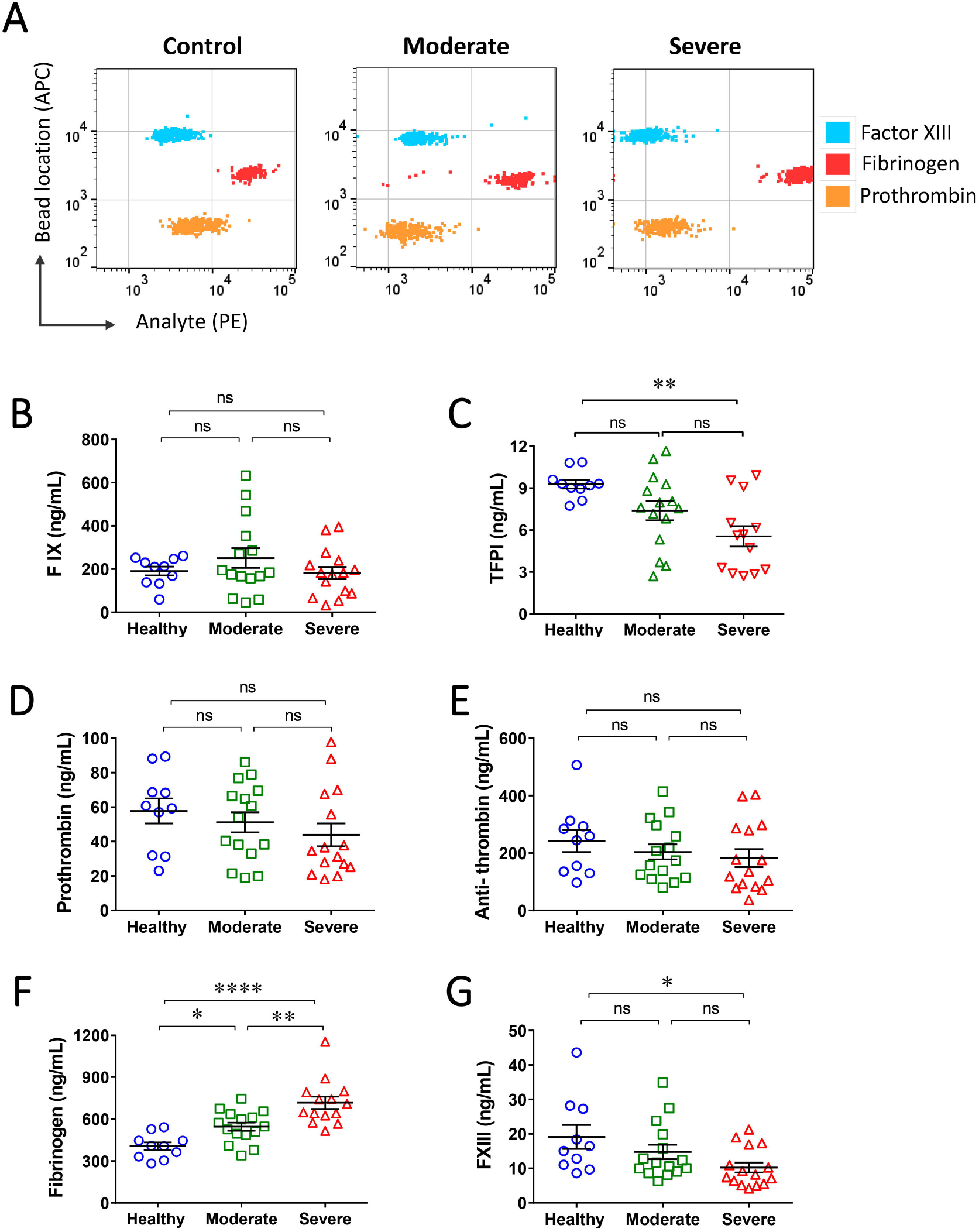
SARS-CoV-2 suppresses TFPI and factor XIII, and induces fibrinogen levels. (**A**) Representative scatter dot plots from flow cytometry-based multiplex assay shows the levels of plasma factor XIII (FXIII), fibrinogen, and prothrombin in COVID-19 patients. (**B**) Analysis shows an unchanged FIX level in COVID-19 cases. (**C**) Scatter dot plot from ELISA shows a significantly low level of tissue factor pathway inhibitor (TFPI) particularly in severe COVID-19 patients vs. healthy controls. Scatter dot plots from multiplex assay show comparable (**D**) prothrombin and (**E**) antithrombin levels, (**F**) significantly higher levels of fibrinogen, and (**G**) lower levels of FXIII in COVID-19 patients vs. healthy controls. *, p<0.05; **, p<0.005; ****, p<0.0001.

### SARS-CoV-2 infection enhances tissue plasminogen activator and fibrinolysis

Aberrant fibrinolysis and elevated levels of D-dimer are commonly seen in COVID-19 patients particularly those who required intensive care. However, it is not clear whether elevated D-dimer is contributed due to dysregulation of fibrinolytic components during the early or late stage of COVID-19 pathogenesis. In this series, the key components of fibrinolysis pathways were assessed in moderate and severe patients. Interestingly, the PAI-1 level was profoundly elevated in the moderate patients and a comparatively lower level was observed in the severe patients (**Fig. 3A-B**). In contrast, the level of tPA in the moderate cases was unchanged however, a significantly higher level was observed in the severe COVID-19 cases (**Fig. 3C**). Moreover, aberrant fibrinolysis was confirmed in the recruited moderate and severe COVDI-19 cases by assessing the D-dimer levels through flow cytometry. Indeed, consistent with the tPA level in plasma, a trend of elevated D-dimer level was observed in the moderate cases and the level was found profoundly elevated in the severe COVID-19 (**Fig. 3D**). Importantly, the plasminogen level was comparable in moderate and severe patients vs. healthy controls (**Fig. 3E**). Overall, these findings strongly suggest that the increased level of PAI-1 in moderate cases likely causes increased resistance to fibrinolysis, which in turn, contributes to thrombosis. However, elevated levels of tPA, particularly in severe COVID-19 cases, promote hyperfibrinolysis and increase the level of D-dimer formation.

**Figure 3:**
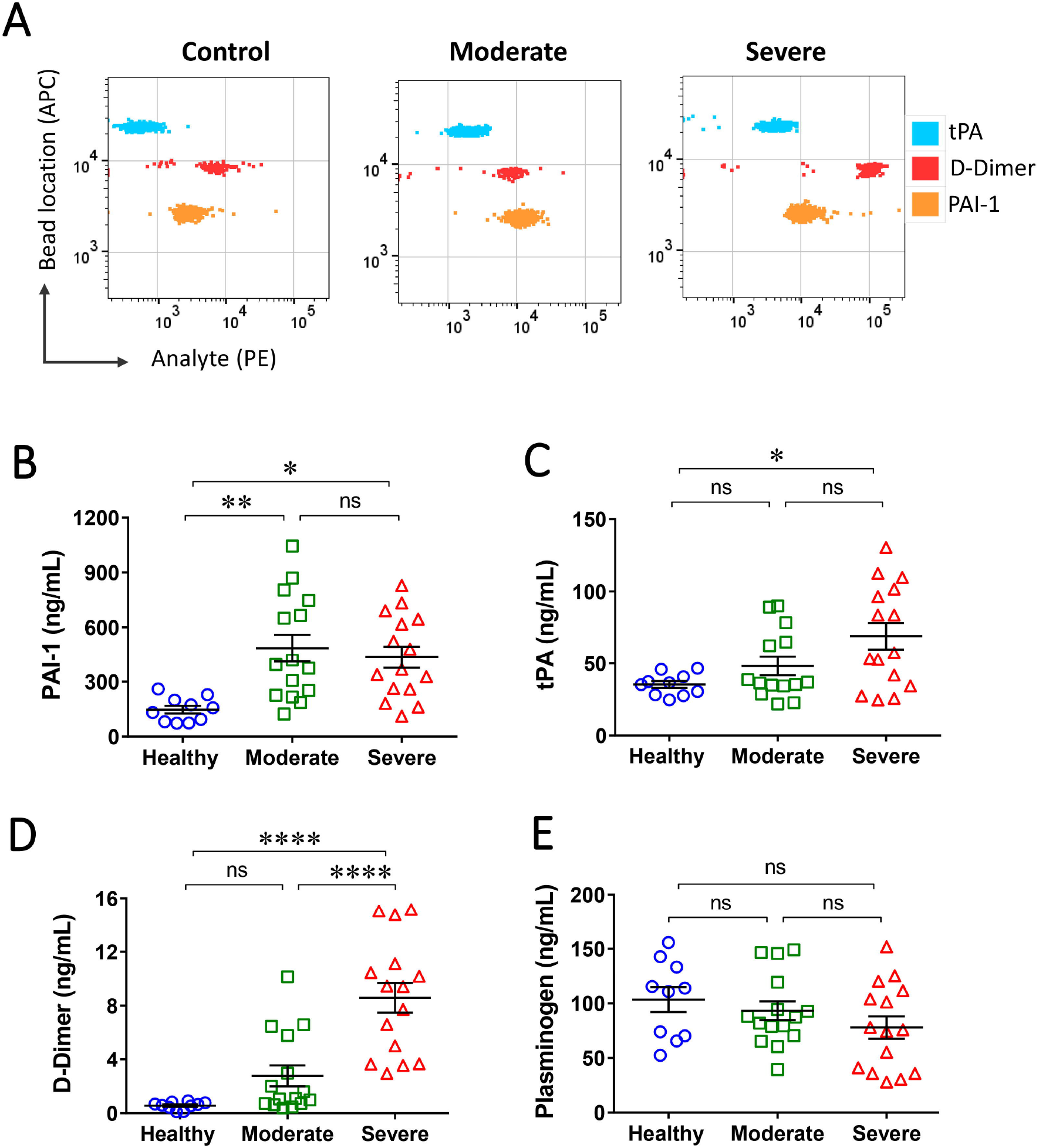
Defective fibrinolysis in COVID-19. (**A**) Representative scatter dot plots from multiplex assay show a higher level of D-dimer, plasminogen activator inhibitor-1 (PAI-1), and tissue plasminogen activator (tPA) in COVID-19 patients. Scatter dot plots show a higher level of (**B**) PAI-1 both in moderate and severe, (**C**) tPA in severe, (**D**) D-dimer particularly in severe COVID-19 vs. healthy controls, and (**E**) comparable plasminogen level between controls and COVID-19 patients. *, p<0.05; **, p<0.005; ****, p<0.0001.

## Discussion

Our findings show profoundly elevated levels of endothelial cell and platelet activation markers P-selectin and sCD40L in moderate cases and the levels were significantly abolished in critically ill patients. Moreover, attenuated levels of TFPI and FXIII, and a higher level of fibrinogen were observed particularly in the severe COVID-19. The level of PAI-1 was elevated in both the patient groups and increased tPA level was seen only in severe COVID-19 cases.

PT and APTT are the laboratory tools that predict the defective extrinsic and intrinsic coagulation pathways.^30^ In contrast to our observations, a study has reported prolonged PT and APTT^31^, while others have shown shortened PT and APTT in severe vs. moderate patients.^32-33^ Consistent with our findings, studies have also reported a comparable PT and APTT between moderate and severe patients.^34-35^ Other coagulation parameters like D-dimer and fibrinogen are consistently reported to be elevated^7,35-36^, however, there are contradictory reports on thrombocytopenia in critical COVID-19 patients.^4,35,37-38^ Therefore, these initial diagnostic tools may not provide a clear clinical scenario of COVID-19 patients and a deeper investigation of coagulation pathway and molecular markers related to platelet activation might help in better clinical interpretation.

Soluble P-selectin, secreted by activated endothelial cells and platelets, not only activates platelets but also act as a key inflammatory mediator during viral infections including influenza ^39-40^ and SARS-CoV-2.^41^ In our study, a higher level of plasma P-selectin in moderate COVID-19 cases, suggests that SARS-CoV-2 infection potentially triggers the release of P-selectin from endothelial cells which eventually contributes to inflammatory reaction and thrombotic complications, in turn, increases the severity of COVID-19. Consistent with P-selectin levels, sCD40L level was profoundly elevated in moderate COVID-19 patients. Platelet-derived sCD40L also provides vital signals to induce B cells activation and immunoglobulin secretion.^42^ Another study highlights the influence of platelet-T-cell interactions during SARS-CoV-2 infections by modulating cytokine IL-17 and IFN-γ production.^43-44^ Moreover, platelet-T cell interaction is clinically relevant as sCD40L was reported to enhance CD8+ T cell activity.^45^ sCD40L plays a larger role in shaping innate immune responses through its cognate receptor, CD40, which is expressed on neutrophils^46^, endothelial cells^47^, and platelets.^48^ This suggests the possible interaction between platelet and neutrophil in moderate COVID-19 which likely plays a role in disease severity. The platelet-neutrophil interaction was also previously reported in various other pathological conditions.^46^ Such interactions in COVID-19 may lead to platelet-mediated neutrophil activation through sCD40L. Moreover, an increased level of sCD40L was found to be associated with plasma CRP level and platelet-monocytes interaction.^49^ These studies signify the contribution of sCD40L in the SARS-CoV-2 -induced pathogenesis.

A trend of lower plasma TFPI levels was observed in the moderate cases and the levels further decreased in the severe COVID-19. The lower level of TFPI particularly in severe patients is likely insufficient to inhibit the procoagulant response which is induced by SARS-CoV-2 –mediated endothelial dysfunction. TFPI deficiency prompts thrombin generation via TF pathway and FX activation that ultimately induces platelet activation (**Fig. 4**). In contrast to our findings, a recent study reported a higher level of TFPI in severe COVID-19 patients which was likely contributed due to heparin-induced TFPI release from endothelial cells as, unlike to our recruited cohort, patients were treated with heparin.^23^ Local TF-dependent coagulation pathway activation and concomitant inhibition of fibrinolysis as indicated by the increased level of PAI-1 particularly in moderate COVID-19 cases are clinical features of ARDS. Such coagulation activation leads to alveolar fibrin deposition which is a pathological feature of early-phase acute lung injury.^50^ It is evidenced that recombinant TFPI (recTFPI) has antithrombotic effects in a variety of *ex vivo* and *in vivo* experimental models ^51^. Pre-clinical studies have evidenced that infusing a large dose of the recTFPI limits disseminated intravascular coagulation (DIC) induced by TF.^52-53^

**Figure 4:**
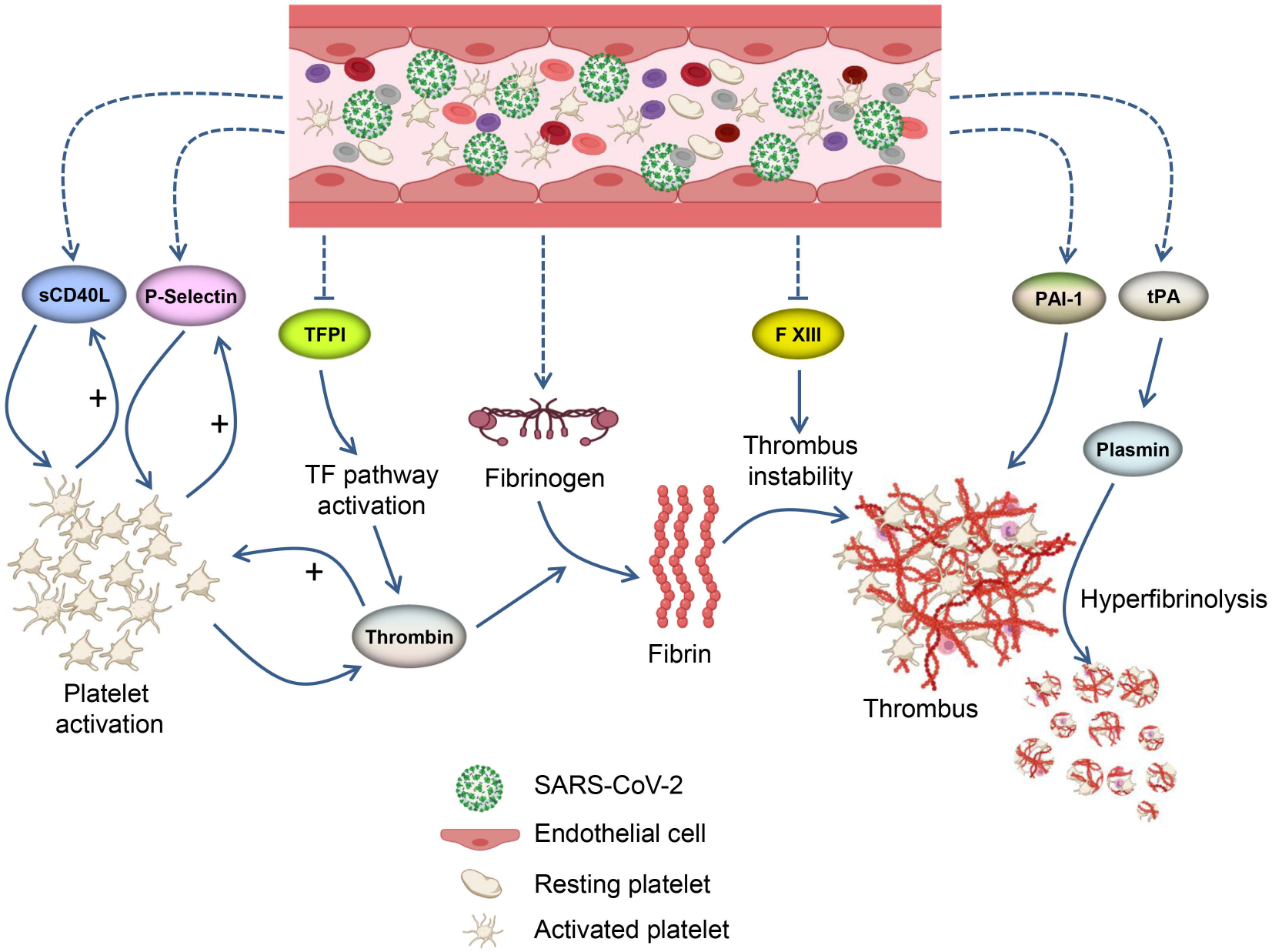
Diagram illustrating the SARS-CoV-2 -mediated induction of plasma CD40L and P-selectin levels, and suppression of tissue factor pathway and FXIII. SARS-CoV-2 -mediated endothelium damage potentially contributes to the elevated level of plasma CD40L (sCD40L) and P-selectin which induces platelet activation. The activated platelet further releases P-selectin and sCD40L and provides positive feedback to platelet plug formation and platelet-mediated thrombin generation. An attenuated level of TFPI activates the tissue factor pathway which likely boosts thrombin level and enhances platelet activation. The elevated level of fibrinogen in COVID-19 patients likely provides the required fibrin however, a decreased level of FXIII indicates the defective fibrin polymerization which potentially results in unstable thrombi formation and embolization. Moreover, an elevated level of PAI-1 prevents fibrinolysis in moderate COVID-19, however, an increased level of tPA in severe COVID-19 likely induces hyperfibrinolysis in turn increased level of D-dimer formation. s**CD40L**; soluble CD40 ligand, **TFPI**; tissue factor pathway inhibitor, **FXIII**; factor XIII, **PAI-1**; plasminogen activator inhibitor-1, **tPA**; tissue plasminogen activator, **+**; positive feedback. Dotted lines; not known if direct or indirect regulation, Solid line; direct regulation.

Our studies revealed a significantly decreased level of FXIII in the severe COVID-19 cases. Emerging pieces of evidence suggest that FXIII crosslink fibrin and promote thrombus stability during platelet accumulation and thrombosis.^54^ Studies employing a ferric chloride-induced venous thrombosis model have shown an increased thrombus embolization in FXIII deficient mice.^55^ Interestingly, FXIII supplementation stabilizes deep vein thrombi in mice and limits pulmonary embolism.^56^ These findings suggest that acquired FXIII deficiency in the severe COVID-19 patients probably causes thrombus instability and embolization. Observed fibrinogen induction in the COVID-19 probably resulted due, partly, to the enhanced inflammatory reaction. It is also evident that fibrinogen and FXIII interact during thrombogenesis to activate FXIII and facilitate the delivery of FXIII to the growing thrombus.^54^ A recent report suggests that FXIII plays a critical role in extravascular fibrinogen crosslinking.^57^ Therefore, further investigation is required to establish if there is any direct correlation between enhanced fibrinogen and decreased FXIII levels in the severe COVID-19.

Aberrant fibrinolysis in COVID-19 patients is consistently reported by multiple studies.^9,37,58-59^ The induction of PAI-1 observed in our COVID-19 patients indicates resistance to thrombus dissolution, at a higher level in the moderate stage which potentially promotes thrombosis. Moreover, elevated levels of tPA particularly in severe COVID-19 indicate the induction of hyperfibrinolysis. The findings are well-correlated with the significantly elevated level of D-dimer in the severe but not in moderate patients. Similar to our finding, recent studies also reported an elevated level of PIA-1 and tPA in hospitalized COVID-19 patients.^24,60^ The study reported that an extremely elevated level of tPA was significantly associated with higher mortality.^24^ The higher level of tPA is consistent with our findings where we have observed a higher level of tPA in the cohort of critically ill COVID-19 patients.

In conclusion, here we report the induction of platelet activation markers in the moderate stage of COVID-19, and the levels markedly declined with the progression of disease severity. A lower level of plasma TFPI in severe COVID-19 indicates the activation of the TF pathway. We show that the attenuated FXIII level was inversely correlated with the level of fibrinogen as COVID-19 severity progressed. The induction of PAI-1 in the moderate stage of the disease due, partly, to endothelial dysfunction and, tPA and D-dimer in severe conditions suggest a switch of fibrinolysis process with the progression of COVID-19 severity. These findings from moderate and severe disease stages would help design a better thromboprophylaxis strategy to minimize the thromboembolism and severity of COVID-19.

## Acknowledgment

The work was supported by COVID-19 (CoV-0302), Targeted (1801090144-P), and Competitive (1901090162-P) research grants from the University of Sharjah to Firdos Ahmad. This work was also supported by the Cardiovascular Research Group at the University of Sharjah.

## Author contribution

AOA, AMY, and MNJ performed experiments and collected data, AWA, ME, FA helped with sample and clinical data collection, MK and RH performed data interpretation and helped in manuscript writing, and FA design study, acquired funding, supervised the project, performed data analysis and interpretation, wrote and revised the manuscript.

